# A natural, conditional gene drive in plants

**DOI:** 10.1101/519884

**Authors:** Anthony J. Conner, Jeanne M.E. Jacobs

## Abstract

A new class of gene drive in plant populations with herbicide resistance is described; a conditional gene drive that operates following herbicide application. Screening progeny from controlled crosses of *Brassica napus* heterozygous for a dominant allele conferring chlorsulfuron resistance, demonstrated that the herbicide imposes *in planta* gametic selection against pollen and ovules with the recessive allele for herbicide susceptibility, as well as embryonic selection against embryos homozygous for the susceptible allele. We postulate that natural gene drives are common in plant populations and can operate in a conditional manner resulting in non-Mendelian inheritance in response to abiotic and biotic stresses.

---

Gene drives involve biased inheritance of genetic elements or specific alleles from parents to offspring through sexual reproduction^1,2^. A consequence of gene drives is an increased frequency of specific genetic elements or alleles and their accelerated spread throughout populations over successive generations^1–4^. Here we describe a conditional gene drive active in plant populations with herbicide resistance. Plants heterozygous for an allele conferring herbicide resistance at a single locus exhibit normal Mendelian inheritance. However, following application of the herbicide, highly distorted segregation of herbicide resistance occurs among progeny. This represents a new class of gene drive that helps to explain the rapid emergence of herbicide resistance in plant populations^5,6^.

Herbicide resistance has become wide-spread in populations of weeds and is commonly the result of point mutations inherited as single dominant mutations^5,7^. The *Brassica napus* mutant line 30a, derived from seed mutagenesis with ethyl methanesulfonate (EMS), is homozygous at a single locus for a dominant allele conferring resistance to the sulfonylurea herbicide chlorsulfuron^8^. Crossing to a wild-type near-isogenic line susceptible to the herbicide generated progeny heterozygous for sulfonylurea resistance. Upon self-pollination of these heterozygous plants, all progeny segregated in a 3:1 ratio for herbicide-resistant and herbicide-sensitive progeny (Supplementary Table 1) as expected. However, when plants of the same heterozygous status were sprayed with chlorsulfuron, virtually all of the progeny of all plants were herbicide-resistant, irrespective of whether plants were sprayed once (Supplementary Table 2) or repeatedly (Supplementary Table 3). We conclude that the application of chlorsulfuron to *B. napus* plants heterozygous for a chlorsulfuron-resistant allele at a single locus results in a markedly distorted segregation toward chlorsulfuron-resistant progeny (Table 1).

**Table 1.** Inheritance of herbicide resistance is conditional upon herbicide application. Segregation of herbicide resistance among the self-pollinated progeny of *Brassica napus* plants heterozygous for a chlorsulfuron-resistant allele at a single locus. Plants were either not sprayed with chlorsulfuron (None), sprayed once when the plants were in the rosette phase (Once), or sprayed when the plants were in the rosette phase followed by repeated sprays every two weeks until initiation of plant senescence (Repeated). The segregation data are for pooled data of progeny from multiple plants (n = 18 to 23), with Chi-square and P-values for one degree of freedom. Data from the individual progeny are presented in Supplementary Tables 1-3.

| **Herbicide treatment** | **Number of herbicide-resistant progeny** | **Number of herbicide-sensitive progeny** | **Chi-square (and P-value) for 3:1 ratio** |
| --- | --- | --- | --- |
| None | 1402 | 464 | 0.018 (P>0.05) |
| Once | 1588 | 17 | 491 (P< 0.001) |
| Repeated | 728 | 2 | 238 (P< 0.001) |

To explore the underlying reason for the skewed segregation, we applied chlorsulfuron at specific times during development of the heterozygous plants, then assessed the self-pollinated and reciprocal backcross progeny for chlorsulfuron resistance. In the absence of herbicide treatment, the segregation of herbicide-resistant and herbicide-sensitive progeny occurred in the expected 3:1 (self-pollinated) and 1:1 (backcrossed) ratios for standard Mendelian genetics (Table 2). Highly skewed segregation among self-pollinated and reciprocal backcrossed progeny occurred following a single application of chlorsulfuron at any time between young seedlings with 2-3 true leaves up until two weeks after the first siliques (seed pods) had reached full size (Table 2). A reduction in seed number per silique following chlorsulfuron application is indicative of the *in planta* elimination of herbicide-sensitive ovules or abortion of herbicide-sensitive embryos. The skewed segregation when the herbicide was applied after siliques had reached full size (i.e. well after embryo formation) signifies an *in planta* effect on embryo abortion. The application of chlorsulfuron to heterozygous pollen parents in the rosette phase prior to flowering, followed by backcrossing to wild-type plants also resulted in distorted segregation in favour of herbicide-resistant progeny (Table 2). However, the skewed segregation selecting against herbicide-sensitive pollen was not as dramatic as when applying the herbicide to the heterozygous ovule parent (Table 2).

**Table 2.**
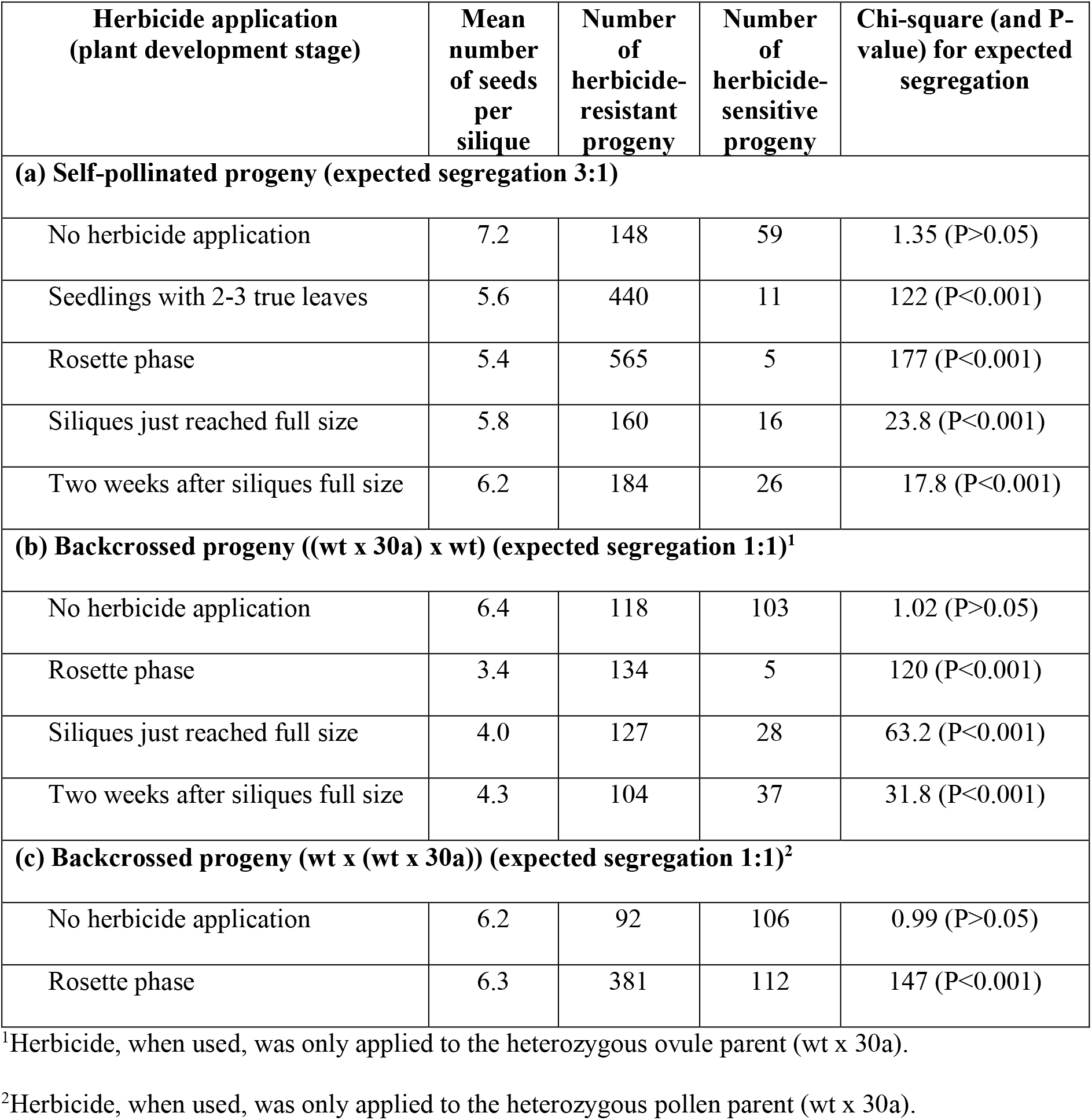
Herbicide application during plant development influences seed set and segregation of herbicide resistance. Segregation of herbicide resistance among the progeny of *Brassica napus* plants (wt x 30a) subjected to chlorsulfuron applied once at different times of plant development. These plants were heterozygous for a chlorsulfuron-resistant allele at a single locus and either self-pollinated (a), backcrossed using pollen from non-sprayed wild type plants (b), or backcrossed as a pollen donor to non-sprayed wild type plants (c). Chi-square and P-values are presented for one degree of freedom.

To distinguish between the elimination of embryos homozygous for herbicide-sensitivity and/or herbicide-sensitive ovules, plants heterozygous for herbicide resistance (wt x 30a) were backcrossed using pollen from non-sprayed plants homozygous for herbicide resistance. As expected, given the homozygous status of the pollen parent, all progeny ((wt x 30a) x 30a) exhibited herbicide resistance irrespective of whether the heterozygous ovule parent was sprayed with chlorsulfuron (n= 111 progeny) or not (n= 223 progeny). However, when chlorsulfuron was applied to the heterozygous ovule parent in the rosette phase prior to flowering the number of seeds per silique was reduced to 3.4, compared with 6.4 when the heterozygous ovule parent was not sprayed. Since all progeny are expected to be herbicide-resistant, no embryo effects are expected. Therefore, the approximate halving of seeds per silique establishes that the *in planta* elimination of the segregating herbicide-sensitive ovules is a major effect contributing to the skewed segregation.

The application of herbicides to plants heterozygous for a herbicide-resistant allele results in a highly biased segregation toward herbicide-resistant progeny (Tables 1 and 2). A series of controlled crosses using pollen and/or ovules from herbicide-treated plants allowed the biased segregation to be attributed to the elimination of segregating herbicide-sensitive gametes and embryos. This will be a consequence of herbicide uptake and translocation throughout the plants, thereby effecting toxicity to pollen, ovule and embryos susceptible to the herbicide. Chlorsulfuron is well known to be rapidly translocated throughout plants in an acropetal manner^9^ and is therefore capable of targeting pollen, ovules and embryos *in planta*. This is supported by sub-lethal foliar applications of sulfonylurea herbicides inducing male sterility in *B. napus* and other cruciferous plants^10–11^. Plants heterozygous for resistance to any systemic herbicide are therefore likely to exhibit similar distorted segregation for herbicide resistance among their progeny following herbicide application. The degree of segregation distortion will be dependent upon the extent of herbicide translocation to segregating pollen, ovules and embryos. Furthermore, the herbicide-resistant alleles need to be expressed in the vegetative tissue, as well as within the reproductive tissue involving gametes and embryos.

This study has described a new class of gene drive active in plant populations with herbicide resistance; a conditional gene drive that only operates upon application of the herbicide. The *in planta* elimination of gametes with a recessive allele for herbicide susceptibility and embryos homozygous for the recessive allele enriches the next generation with individuals homozygous or heterozygous for dominant alleles conferring herbicide resistance. As a consequence the frequency of herbicide-resistant alleles in subsequent generations is increased.

Gene drives have been described and developed in animal systems. Engineered gene drive systems offer approaches to control invertebrate and vertebrate pests^1,3,12,13^, with some theoretical opportunities also proposed for plants^1,13^. Ironically, these proposed gene drives for plants involve scenarios for the elimination of herbicide-resistant biotypes in plant populations, whereas this study clearly demonstrates that natural gene drives can account for the rapid increase of the frequency of herbicide-resistant alleles in natural plant populations. Gene drives are poorly documented and not well understood in plants. Distorted segregation in plants is commonly recognised and is often associated with cytological abnormalities or genetic curiosities^14^. These particular events generally have poor transmission from parents to offspring relative to the wild-type and are usually maintained as genetic stocks with limited occurrence in natural populations. Two examples recognised as gene drives in plants involve sex expression associated with gynodioecy in *Silene acaulis*^15^ and a centromere-associated locus in *Mimulus* interspecific hybrids^16^.

This study has illustrated a simple conditional gene drive likely to operate as a natural process in environments where herbicides are commonly used and where herbicide resistance alleles are present in weed populations. It represents a new class of gene drive that helps to explain the rapid emergence of herbicide resistance in plant populations^5,6^. We postulate that natural gene drives are common in plant populations and can operate in a conditional manner during sexual reproduction to effect a biased inheritance for specific alleles conferring resistance to abiotic and biotic stresses, including xenobiotic chemicals, acid soil, metal toxicity, temperature stress and possibly pathotoxins, viruses and viroids. Effecting a conditional gene drive through herbicide treatments provides an eloquent example to target the induction of biased segregation in favour of a specific allele. This is due to the potent nature of systemic herbicides to translocate acropetally and effectively eliminate herbicide sensitive gametes and embryos *in planta*. Conditional gene drives for other forms of abiotic and biotic stress may exhibit less penetrance and a reduced bias of segregation ratios, but can be still effective at increasing the frequency of specific alleles in subsequent generations.

This new concept of conditional gene drives has important implications for interpreting inheritance of traits in plants. The environmental conditions imposed upon the growth of parental plants can establish a gene drive that effects selection against recessive alleles during sexual reproduction of heterozygous individuals. Screening of subsequent progeny may result in an over-representation of dominant alleles conferring resistance to abiotic and biotic stresses, with some traits being more simply inherited than otherwise apparent. Further development of conditional gene drives may offer mitigation for the concerns raised about unintended impacts associated with the introduction of engineered gene drive systems to control pests, weeds and diseases^1,2,4,13,17,18^. Most importantly, this form of gene drive provides a fundamental mechanism for the interpretation of rapid natural selection in adaption of plants to abiotic and biotic stresses in changing environments.

## Methods

Two near-isogenic *Brassica napus* lines were used in this study; a wild-type herbicide-susceptible line (wt) and 30a, an ethyl methanesulfonate-induced mutant homozygous for single dominant allele conferring resistance to the sulfonylurea herbicides^8^. These were derived from a rapid cycling *B. napus* line CrGC#5, originally obtained from the Crucifer Genetics Cooperative, University of Wisconsin. The two lines were hybridised to generate a population heterozygous for herbicide resistance (wt x 30a). Some experiments involved backcrosses of the heterozygous plants to either wt or 30a. Plants were grown individually in 1.8-litre plastic pots (15 cm diameter at top tapering to 10 cm diameter at base, 15 cm high) with a standard potting mix and maintained in a greenhouse as previously described^19^.

When required, plants were sprayed with chlorsulfuron (a sulfonylurea herbicide) at a rate of 3 mg/litre until runoff. This treatment was lethal to the wild-type plants, but had no growth impact on the plants heterozygous or homozygous for chlorsulfuron resistance. Following self-pollination or backcrossing, all seeds were harvested at maturity. Progeny were screened for chlorsulfuron resistance as previously described^20^. Segregation among progeny was assessed by chi-square goodness of fit tests to determine whether the observed segregation deviated from the expected 3:1 or 1:1 ratios for Mendelian genetics.

### Data availability

All data generated or analysed during this study are included in this published article (and its supplementary information files).

## Acknowledgements

We thank Suzanne Lambie for screening the progeny for herbicide resistance and Peter Dearden for discussions and comments on the manuscript.

## Author Contributions

A.J.C and J.M.E.J conceived and designed the experiments, collated and interpreted the data, and wrote the manuscript. A.J.C grew the plants, applied the treatments, made the crosses and harvested the seed.

## Competing interests

The authors declare no competing interest

