## Supplementary Tables 1-3 for "A natural, conditional gene drive in plants"

**Supplementary Table 1**

Segregation of herbicide resistance among the self-pollinated progeny of *Brassica napus* plants not sprayed with chlorsulfuron. The plants were heterozygous for a chlorsulfuron-resistant allele at a single locus and expected to segregate in a 3:1 ratio for herbicide-resistant and herbicide-sensitive individuals among the progeny.

| Plant number (wt x 30a) | Number of herbicide-resistant progeny | Number of herbicide-sensitive progeny | Chi-square for 3:1 ratio[1] |
|---|---|---|---|
| i | 61 | 20 | 0.00 |
| ii | 46 | 16 | 0.02 |
| iii | 53 | 16 | 0.12 |
| iv | 81 | 28 | 0.03 |
| v | 48 | 23 | 2.07 |
| vi | 62 | 27 | 1.35 |
| vii | 79 | 29 | 0.20 |
| viii | 54 | 14 | 0.71 |
| ix | 67 | 14 | 2.57 |
| x | 39 | 21 | 3.20 |
| xi | 47 | 12 | 0.68 |
| xii | 53 | 19 | 0.07 |
| xiii | 93 | 30 | 0.02 |
| ixv | 80 | 21 | 0.95 |
| xv | 66 | 22 | 0.00 |
| xvi | 48 | 20 | 0.71 |
| xvii | 108 | 37 | 0.02 |
| xviii | 3 | 3 | 2.00 |
| ixx | 69 | 19 | 0.55 |
| xx | 22 | 12 | 1.92 |
| xxi | 109 | 22 | 4.71 |
| xxii | 65 | 16 | 1.19 |
| xxiii | 49 | 23 | 1.85 |

[1]Chi-square values less than 3.84 for one degree of freedom are indicative of the observed segregation being not significantly different at the 5% probability level from the expected 3:1 segregation ratio for herbicide-resistant and herbicide-sensitive progeny.

**Supplementary Table 2**

Segregation of herbicide resistance among the self-pollinated progeny of *Brassica napus* plants sprayed once with chlorsulfuron just as the plants were beginning to bolt to flower formation. The plants were heterozygous for a chlorsulfuron-resistant allele at a single locus and expected to segregate in a 3:1 ratio for herbicide-resistant and herbicide-sensitive individuals among the progeny.

| Plant number (wt x 30a) | Number of herbicide-resistant progeny | Number of herbicide-sensitive progeny | Chi-square for 3:1 ratio[1] |
|---|---|---|---|
| i | 4 | 0 | 1.33 |
| ii | 125 | 0 | 41.7 |
| iii | 83 | 7 | 14.2 |
| iv | 73 | 1 | 22.1 |
| v | 75 | 0 | 25.0 |
| vi | 105 | 0 | 35.0 |
| vii | 125 | 0 | 41.7 |
| viii | 67 | 0 | 22.3 |
| ix | 86 | 1 | 26.4 |
| x | 71 | 1 | 21.4 |
| xi | 89 | 2 | 25.2 |
| xii | 88 | 0 | 29.3 |
| xiii | 94 | 1 | 29.1 |
| ixv | 56 | 0 | 18.7 |
| xv | 80 | 1 | 24.4 |
| xvi | 76 | 0 | 25.3 |
| xvii | 145 | 2 | 43.8 |
| xviii | 92 | 0 | 30.7 |
| ixx | 54 | 1 | 15.8 |

[1]Chi-square values greater than 3.84 for one degree of freedom are indicative of the observed segregation being significantly different at the 5% probability level (i.e. distorted segregation) from the expected 3:1 segregation ratio for herbicide-resistant and herbicide-sensitive progeny.

**Supplementary Table 3**

Segregation of herbicide resistance among the self-pollinated progeny of *Brassica napus* plants sprayed with chlorsulfuron just as the plants were beginning to bolt to flower formation, and then repeatedly every two weeks until initiation of plant senescence. The plants were heterozygous for a chlorsulfuron-resistant allele at a single locus and expected to segregate in a 3:1 ratio for herbicide-resistant and herbicide-sensitive individuals among the progeny.

| **Plant number (wt x 30a)** | **Number of herbicide-resistant progeny** | **Number of herbicide-sensitive progeny** | **Chi-square for 3:1 ratio[1]** |
|---|---|---|---|
| i | 5 | 0 | 1.67 |
| ii | 50 | 0 | 16.7 |
| iii | 78 | 0 | 26.0 |
| iv | 50 | 0 | 16.7 |
| v | 48 | 0 | 16.0 |
| vi | 83 | 0 | 27.7 |
| vii | 52 | 0 | 17.3 |
| viii | 30 | 1 | 7.84 |
| ix | 20 | 0 | 6.67 |
| x | 10 | 0 | 3.33 |
| xi | 40 | 0 | 13.3 |
| xii | 60 | 0 | 20.0 |
| xiii | 17 | 0 | 5.67 |
| ixv | 10 | 0 | 3.33 |
| xv | 57 | 1 | 16.8 |
| xvi | 100 | 0 | 33.3 |
| xvii | 37 | 0 | 12.3 |
| xviii | 7 | 0 | 2.33 |

[1]Chi-square values greater than 3.84 for one degree of freedom are indicative of the observed segregation being significantly different at the 5% probability level (i.e. distorted segregation) from the expected 3:1 segregation ratio for herbicide-resistant and herbicide-sensitive progeny.
